# Effects of Willow and Pine tree growth in bacterial abundance, interactions and metabolic profile in a copper mine tailing

**DOI:** 10.1101/2025.04.28.651128

**Authors:** Jaime Ortega, Gabriel Gálvez, Gladis Serrano, Jorge Torres, Víctor Aliaga-Tobar, Emilio Vilches, Angélica Reyes, Vinicius Maracaja-Coutinho, Valentina Parra, Alex Di Genova, Lorena Pizarro, Mauricio Latorre

## Abstract

The Cauquenes tailings is one of the largest mine tailings in Chile and represents a significant environmental hazard due to its high copper levels and acidic pH. However, there is evidence of willow and pine tree growth in these terrains, which is unexpected given the extreme conditions of this environment. Considering the importance of plants in restoration strategies, the objective of this study was to analyze the effect that willow and pine growth have on the heavy metal contents of their surrounding soil, and the abundance, interaction, metabolic profile and assembly of the bacterial community. In the Cauquenes tailing, we collected samples from willow and pine vegetated soils as well as the surrounding non-vegetated soils. We analyzed characteristics such as heavy metal concentrations and pH levels in all sampled soils, and in tree leaves and roots. Vegetated sites exhibited lower copper levels and a neutral pH, while both tree species accumulated heavy metals in their tissues. We also determined bacterial abundance in all soil samples, and co-occurrence networks were constructed to assess bacterial interactions. Both trees promoted a higher abundance of Proteobacteria and had networks with higher modularity compared to non-vegetated soils. Finally, we estimated the metabolic profile of the bacterial community using PICRUSt 2 analysis and determined carbon source utilization with BioLog Ecoplates. PICRUSt 2 predicted higher metabolic activity in vegetated soils, and BioLog Ecoplate analysis revealed the utilization of a greater number of carbon sources. The microbial communities from vegetated soils were assembled in a deterministic manner, while the community from non-vegetated soils were stochastically assembled. In conclusion, tree growth mitigated the extreme conditions of the tailing and promoted a more active and healthier bacterial community.

**Highlights:**

- Willow and pine tree can grow and mitigate toxic metal conditions in the tailings
- Both trees provoked a shift in bacterial abundance and interaction network
- The provoked shift was tree-dependent
- Microbial activity was higher in vegetated soils

## 1. Introduction

Mining activities, while important for the economic growth and development of industrialized and developing countries (Rashed, 2010), are responsible for the accumulation of potentially toxic elements in the form of tailings, which pose a high environmental risk (Buch et al., 2021). These tailings can contaminate the surrounding soil systems and inland water bodies via heavy metal (HMs) dispersion, altering the soil microorganisms, plants and animals (Wen et al., 2024). Chile is the country with the largest number of copper tailings worldwide. In particular, the Cauquenes tailings deposit is one of the oldest and largest derived from an underground mine (El Teniente mine). Covering a total area of 12.5 km^2^, Cauquenes tailing has a high amount of copper (Cu) (concentration over 1,6 g/L of soluble Cu) and acidic pH of 3 (Galvez et al., 2022), which implies extreme adverse conditions for the development of various organisms.

Copper toxicity in plants impacts various processes, including root and shoot length, nutrient distribution, damage from reactive oxygen species (ROS), photosynthetic performance, and nutrient uptake (Lequeux et al., 2010; Yang et al., 2002). Specifically, root shortening exposes plants to water stress, which further hinders their growth in this type of soil (Guittonny-Larchevêque et al., 2016). In turns, low pH exacerbates these copper toxic effects, causing deficiencies of essential nutrient such as nitrogen, phosphorus, potassium, and calcium, while simultaneously increasing the presence of toxic elements like aluminum, manganese, and iron in the soil (Zhao et al., 2014).

Despite the presence of these toxic effects, the natural growth of different plants in copper contaminated environments can be observed. Närhi et al. investigated the plant composition in a tailing-contaminated wetland, revealing that *Carex rostrata*, *Equisetum palustre*, and *Eriophorum angustifolium*, tolerated high concentrations of soil Cu, Zn, among other HM (Närhi et al., 2012). Espinoza et al. analyzed HMs concentrations in sclerophyllous vegetation from the surrounding soils of Cauquenes tailings, finding that *Acacia caven* and *Quillaja saponaria* could accumulate Cu and Zn in their leaves and stems, respectively (Espinoza et al., 2022). These studies suggest that certain factors enable this growth. Within them, interactions between plants and bacterial communities are fundamental for growth, development, and plant health (Abdelaal et al., 2021), since bacterial communities play a crucial role in ecological processes such as cycling of nutrient elements (Yin et al., 2023).

In regard to microbial communities in heavy-metal contaminated environments, it has been shown that they could reduce the toxicity and mobility of heavy metals by altering its forms or their speciation (Kang et al., 2020; Luo et al., 2020; X. Sun et al., 2020). These communities can be sensitive to heavy metals, with copper often causing shifts in their composition and structure. High copper concentrations reduce microbial diversity while favoring resistant taxa like *Pseudomonas, Acinetobacter,* and *Serratia* (Turpeinen et al., 2004).

The presence of plants significantly alters these microbial dynamics. Wang et al. showed that the rhizosphere of copper-tolerant plants, like *Elsholtzia splendens*, exhibits higher microbial biomass and diversity compared to unplanted soils, driven by plant-microbe interactions (Y. P. Wang et al., 2008). Furthermore, Pérez-de-Mora et al. highlighted that plant presence not only influences microbial biomass but also fosters distinct microbial community structures, driven by root exudates and interactions within the rhizosphere (Pérez-De-Mora et al., 2006). Long-term exposure to copper also reveals intriguing patterns; Shaw et al. found that high copper concentrations lead to lower beta diversity while maintaining or even increasing alpha diversity, suggesting selection for specialized microbial taxa (Shaw et al., 2020). Collectively, these studies underscore that the interplay between plants and microbial communities profoundly affects bacterial abundance, interactions, and metabolic profiles in heavy-metal-contaminated environments, such as copper mine tailings.

Understanding the complex microbial relationships within the soil ecosystem requires not only identifying the microbial species present but also exploring how they interact with each other and the environment (C. Gao et al., 2022). To achieve this, co-occurrence networks can be employed. These networks graphically represent potential symbiotic, competitive, and antagonistic relationships by analyzing which microbes frequently appear together across different samples (Peschel et al., 2021a). Studies on metal-contaminated soils have shown that heavy metals significantly alter microbial network complexity, often increasing positive associations among metal-tolerant taxa to maintain community stability (C. Sun et al., 2022; Xu et al., 2023). In copper mine tailings, network analyses reveal distinct interaction patterns, with a shift from simpler networks in low-contamination areas to highly connected networks dominated by generalist taxa in areas with higher contamination (Q. Qi et al., 2022). Additionally, the presence of plants can enhance microbial network complexity by increasing nutrient availability through root exudates, fostering more intricate interactions and greater ecological resilience (C. Sun et al., 2022). The assembly of these microbial communities, shaped by both deterministic and stochastic processes, is influenced by environmental factors like heavy metal concentrations and the presence of vegetation, with deterministic processes often dominating in these highly selective environments (Stegen et al., 2013; Y. Wang et al., 2024; Yin et al., 2023). This approach provides a valuable framework for unraveling the ecological processes governing microbial interactions in contaminated environments.

Given the observed growth of pine and willow in the harsh conditions of Cauquenes tailings, soil samples were taken near and far from those trees. Each sampled was sequenced, Amplicon Sequence Variants (ASVs) were assigned and co-ocurrence networks were constructed to compare bacterial community interactions at each site. This strategy permitted to decipher specific ecological interactions that underpin the resilience and adaptation strategies of pine and willow in metal-contaminated environments, identifying key microbial taxa that associate with these two different pioneer plants, revealing particular bacterial community structures that are essential for survival and growth in such hostile conditions. To further understand the processes governing these communities, we performed a Beta Nearest Taxon Index (βNTI) analysis to determine whether the bacterial assembly is predominantly stochastic or deterministic (Yin et al., 2023). By correlating βNTI values with the presence of vegetation, we seek to elucidate if pine and willow impose deterministic pressures on microbial community assembly through environmental filtering or whether stochastic processes, such as dispersal limitation, dominate in non-vegetated soils. Through this approach, we aim to explore how the presence of pioneer plants configures both the structure and functional potential of microbial communities, shedding light on the ecological dynamics within these extreme environments and their implications for bioremediation strategies

## 2. Materials and methods

### 2.1 Sample collection

Soil samples (∼250 g each) were collected in January 2020 (austral summer) under sterile conditions from the upper 5 to 10 cm of the soil surface at four distinctive zones: the first zone corresponds to the soil close (approximately 10 cm from the root) to willows within the tailings impoundment, hereafter referred to as Willow vegetated soil (WVS) (3 total replicates); the second zone belongs to the soil close to pines within the tailings impoundment, referred to as Pine vegetated soil (PVS) (3 total replicates); the third zone represents the comparative site of site 1 without plant intervention, far site of the willow (approximately 5 meters from the root), referred to as Willow non-vegetated soil (WNVS) (3 total replicates) and the fourth zone represents the comparative site of site 2 without plant intervention, far site of the pine (approximately 5 meters from the root), referred to as Pine non-vegetated soil (PNVS) (3 total replicates). For a total of 12 total samples. From each replicate, a subsample of 200 g was immediately stored on dry ice for microbial analyses (stored for approximately one week), while another subsample of 50 g was air-dried, sieved (≤ 2mm) and stored for physicochemical analyses.

### 2.2 Soil DNA extraction

DNA was extracted from subsamples collected from each soil samples at each site using the Qiagen kit DNeasy Blood & Tissue, combining the manufacturer’s instructions and the CTAB based method (Zhou et al., 1996). Five grams of soil were resuspended in 5 ml extraction buffer [100 mM Tris-HCl; pH 8, 100 mM Na EDTA; pH 8, 100 mM Na2HPO4, 1.5 M NaCl, 1% (w/v) CTAB] and then 25 µl of proteinase K was added and mixed by vortex, followed by incubation at 65 °C for 2 h, with constant mixing. The mixture was centrifuged at 4000×g for 10 min at room temperature and the supernatant fuid was transferred to a clean tube, to which 0.5 volume of ethanol 100% was added. Then, the samples were vortexed for 10 seconds and the mixture was transferred into a DNeasy mini spin column to continue the kit protocol. Concentrations of DNA were evaluated with a nanodrop. Microbial DNA was amplified using a bacteria-specific primer set, 28 F (5′-GA GTT TGA TCM TGG CTC AG-3′) and 519 R (5′-GWA TTA CCG CGG CKG CTG-3′), flanking variable regions V1-V3 of the 16 S rRNA gene (Turner et al., 1999), with barcode on the forward primer. Amplification was performed using the promega GoTaq® G2 Flexi DNA Polymerase Mix, under the following conditions: initial denaturation at 94 °C for 3 min, followed by 28 cycles, each set at 94 °C for 30 seconds, 53 °C for 40 seconds and 72 °C for 1 min, with a final elongation step at 72 °C for 5 min. After amplification, PCR products were checked in 2% agarose gels to determine the success of the amplification and then stored at −20 °C until DNA analyses.

### 2.3 Soil physicochemical measurements

For the measurement of nutrients and pH, 5 grams of sample were placed in a sterile 15 ml tube; then 1:1 (w/v) distilled water was added (Thomas & Wimpenny, 2012). The samples were homogenized for 2 hours at room temperature and then centrifuged at 12000g for 5 minutes. The soluble fraction was recovered. The pH quantification was performed directly from the soluble fraction. Nutrient measurement was performed using 500 uL of the soluble fraction using the Total Reflection X-Ray Fluorescence (TXRF) spectroscopic technique, following a previously described protocol (Tapia et al., 2003).

### 2.4 16S rRNA gene amplification and sequencing

Each sample was sequenced in triplicate (9 samples per site, 36 samples in total), each sample was sent to the Mr. DNA company for amplification and subsequent sequencing of the 16S rRNA gene, using the illumina® MiSeq kit. This service was performed by sequencing the V3 and V4 hypervariability region. The sequences were processed following previously described protocols (Handl et al., 2011). The sequences were overlapped and grouped by samples, to later eliminate the “barcode” segments of the sequences. Sequences less than 150 bp or with ambiguous base allocation were not considered for further analysis. The sequences accepted as valid were grouped using the UClust algorithm (v2.22) with 4%, to eliminate chimeras and groups with a single sequence (singletons) (Edgar, 2010). After being processed, the sequences obtained were analyzed with the Quantitative Insights Into Microbial Ecology 2 (QIIME 2) Program (Caporaso et al., 2010), using the “qiime feature-classifier classify-sklearn” protocol. The sequences were then compared against Greengenes databases (McDonald et al., 2012), using 97% similarity, directly assigning the taxonomy from the closest match. To assign the ASVs, they were filtered by a minimum of 4 reads (readings) per sample, eliminating those ASVs corresponding to mitochondria, chloroplasts and not classified within the bacterial range. The rarefaction analysis was performed using the “qiime diversity alpha-rarefaction” code, setting the number of readings at the lowest obtained from the samples. To obtain the alpha diversity the Shannon and evenness index were calculated, using the code “qiime diversity core metrics phylogenetic”. To construct the phylogenetic tree, we generated a tree file from the previously taxa obtained using the “qiime phylogeny align-to-tree-mafft-fasttree” code and visualized the associations using the ITOL platform (Letunic & Bork, 2021). To predict the supposed metagenomic profile, based on ASV abundance, Phylogenetic Investigation of Communities by Reconstruction of Unobserved States (PICRUSt2 https://github.com/picrust/picrust2; Version 2.5.1) was applied after the processing of amplicon sequencing analysis of the sample. To analyze the PICRUSt 2 results the database MetaCyc was used.

### 2.5 Construction of co-occurrence networks and data analysis

The networks were constructed with the abundance tables obtained from the processing and analysis of bacterial sequences. Co-occurrence network analysis was performed using the R package NetCoMi (Peschel et al., 2021b). Spearman correlations were calculated between taxa, multiplicative replacement (multRepl) was used for replacing zero values and a centered log-ratio transformation (clr) was used for normalization. A threshold of 0.6 was used to select edge connections between taxa pairs. The netAnalyze function was used to calculate network properties and centralities for all nodes were computed. Measurements of the degree, betweenness, closeness, and eigenvector were normalized. The nodes with the highest eigenvector value were considered network hubs. Comparative analyses between the constructed networks were carried out using the netconfer website (Nagpal et al., 2020). To determine the community assembly process, βNTI (Beta nearest taxon index) was calculated for each sampling site as previously described (Stegen et al., 2012). The βNTI metric quantifies the deviation of observed phylogenetic turnover from null expectations, with |βNTI| > 2 indicating deterministic processes (homogeneous selection for βNTI < −2 and variable selection for βNTI > 2) and |βNTI| < 2 indicating stochastic processes. The analysis was performed using the R package “picante”. This analysis was carried out for all samples, and the results were compared across sites with and without tree growth to evaluate the influence of vegetation on community assembly mechanisms.

### 2.6. Determination of carbohydrate consumption

Biolog EcoPlates (Biolog, Inc., Hayward, CA) were used to assess the metabolic activity of microorganisms present in the different soils samples based on the consumption of different carbon sources. Biolog EcoPlates are composed of 31 different carbon compounds and a control. It contains three replicates of the carbon source and control wells. A redox dye (tetrazolium violet) is added in each well, which turns purple when the carbon source is used by the microbial communities present in the sample. Soil samples were mixed in 1:1 w/v with PBS 1X and mixed thoroughly for 1 hour. After that, each well of EcoPlates plate was inoculated with 130 μL of the soil suspension and incubated at 25°C. Purple color development was measured with an HTX microplate reader at 590 nm. This measurement was carried out daily for up to ten days, in which all plates showed color development of at least 1 well.

## 3. Results

### 3.1 Physicochemical Characteristics of the Tailing

The Cauquenes tailing is divided into various sectors based on the concentration of salts, metals, and pH, which are related to the mining activity timeline (Galvez et al., 2022). Interestingly, natural tree growth has been observed in two distinct patches within the area, one with pines and the other with willows (Figure 1). To assess the effect of tree growth on soil concentrations of salts, metals, and pH, samples were collected near willow growth (WVS), pine growth (PVS), and from areas distant from willows (WNVS) and pines (PNVS) (Table 1). The data indicates significant differences in the amounts of metals and salts at these sites. Specifically, WVS exhibited higher concentrations of P, K, and Ca, while pH was neutral, and concentrations of Cu, Na, Zn, and Fe were lower compared to WNVS. Similarly, in PVS, there were higher concentrations of P, K, and Ca, with lower levels of Cu, Fe, and Mn, and a neutral pH compared to PNVS. Notably, both areas with tree growth showed lower Cu concentrations, suggesting a potential phytoremediative or phystosabilizing effect of the trees.

**Figure 1.**
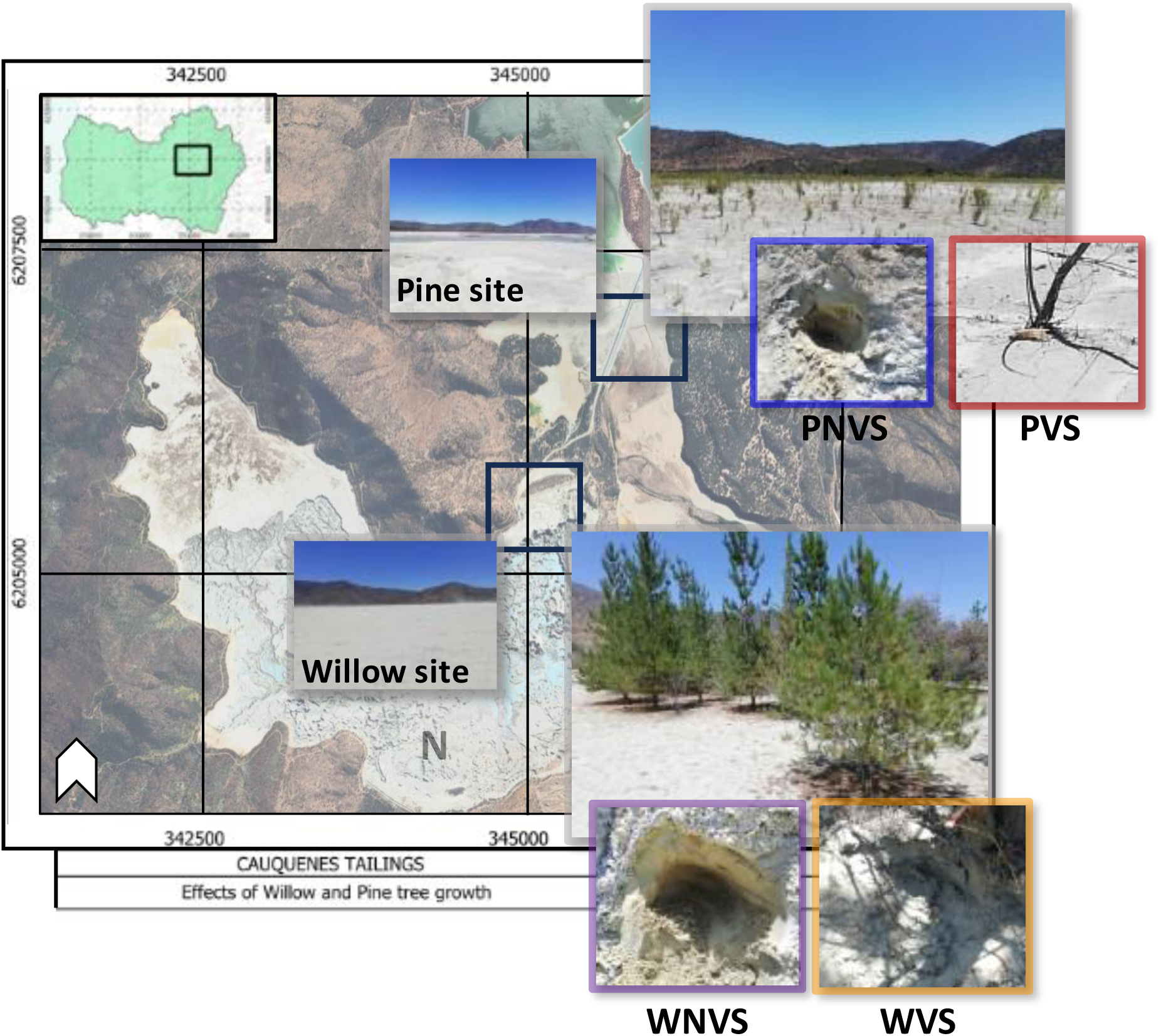
Geographical location of Cauquenes tailing and sampling sites. Blue color indicates Pine Non-Vegetated Soil (PNVS); Red indicates Pine Vegetated Soil (PVS); Purple indicates Willow Non-Vegetated Soils, and Orange indicates Willow Vegetated Soil (WVS).

**Table 1.** Physical and chemical properties of each sector of the Cauquenes tailings. WVS: Willow-Vegetated Soil; WNVS: Willow Non-Vegetated Soil; PVS: Pine-Vegetated Soil; PNVS: Pine Non-Vegetated Soil.

| Element (µg/L) | WVS | WNVS | PVS | PNVS |
| --- | --- | --- | --- | --- |
| Ca | 75416.8 ± 9888.2 | 43918.9 ± 16307 | 6186.6 ± 3059 | 6112.9 ± 715 |
| Cu | 83 ± 39.9 | 25357.5 ± 5471.2 | 35.4 ± 19.8 | 1546 ± 1061.4 |
| Fe | 529.8 ± 407.2 | 1460.7 ± 719.5 | 246.5 ± 157.6 | 442.2 ± 178 |
| K | 20057.9 ± 2394.6 | 2417.1 ± 603.8 | 4444.9 ± 401.8 | 4037.5 ± 576.3 |
| Mn | 16.8 ± 5.8 | 511.2 ± 137.3 | 3.2 ± 1.8 | 67.9 ± 17.6 |
| Na | 3270.2 ± 1083.4 | 8095.4 ± 2170.3 | 1224.8 ± 667 | 1625.4 ± 1183.8 |
| P | 7868.9 ± 1451.7 | 5292.5 ± 1787.8 | 168.4 ± 86.6 | 48.8 ± NA |
| Zn | NA | 525.1 ± 144.2 | NA | 15.5 ± 5.4 |
| pH (unit) | 6.3 ± 0.1 | 4.1 ± 0.1 | 7.1 ± 0.1 | 4.6 ± 0.1 |

To verify this, leaf and root samples from each tree were analyzed for the same elements (Table 2). For willows, leaf (WL) and root (WR) samples showed higher concentrations of all elements compared to WVS, particularly higher Cu and Fe in WR than WL, indicating possible phytostabilization of these metals. Similarly, pine leaf (PL) and root (PR) samples showed higher concentrations of elements compared to PVS. However, unlike willows, PL had higher concentrations of all elements except Na and P compared to PR. The accumulation of metals in leaves could be useful for phytoextraction, as they can be harvested without removing the entire tree (Shahid et al., 2017).

**Table 2.** Chemical elements in roots and leaves of each tree. WL: Willow Leaves; WR: Willow Roots; PL: Pine Leaves; PR: Pine Roots.

| Element (mg/L) | WL | WR | PL | PR |
| --- | --- | --- | --- | --- |
| Ca | 198.4 ± 30.4 | 125.1 ± 35.4 | 75.3 ± 30.6 | 55.7 ± 22 |
| Cu | 1 ± 0.2 | 2.5 ± 0.5 | 2.9 ± 0 | 0.7 ± 0.3 |
| Fe | 11.1 ± 1.3 | 22 ± 3.1 | 17.3 ± 3.2 | 11 ± 4.8 |
| K | 392.5 ± 201.3 | 308.8 ± 87.5 | 188.4 ± 17.7 | 104.2 ± 9.3 |
| Mn | 18.6 ± 8.6 | 1.9 ± 0.8 | 1.2 ± 0.1 | 0.4 ± 0.1 |
| Na | 200.4 ± 112.2 | 51 ± 21.6 | 11.8 ± 4.1 | 776.1 ± 1539 |
| P | 86.1 ± 5.3 | 62.8 ± 7 | 49 ± 2.1 | 470 ± 1285.9 |
| Zn | 4.8 ± 3.2 | 1.9 ± 1.4 | 0.6 ± 0 | 0.3 ± 0.1 |

### 3.2 Abundance and Diversity

As tree presence also influences soil microbial communities, a 16S gene analysis was performed for the four sectors to determine bacterial abundance and diversity. A total of 226, 1199, 307, and 435 amplicon variant sequences (ASVs) were obtained from WVS, PVS, WNVS, and PNVS samples, respectively, with PVS being the most taxonomically rich site. The identity and relative abundance of ASVs for each site are shown in Figures 2A and 2B. WVS was dominated by ASVs belonging to the phyla *Actinobacteria* and *Proteobacteria*, with relative abundances of 31.3% and 31.1%, respectively. In contrast, *Firmicutes* was the most represented phylum in WNVS, with 26.8% relative abundance, compared to 17% in WVS. Similar differences were observed for *Actinobacteria* and *Proteobacteria* in WNVS, with lower abundances (9.38% and 16%, respectively). These results suggest that willows influence the microbial community composition, differing from sites without these trees.

**Figure 2.**
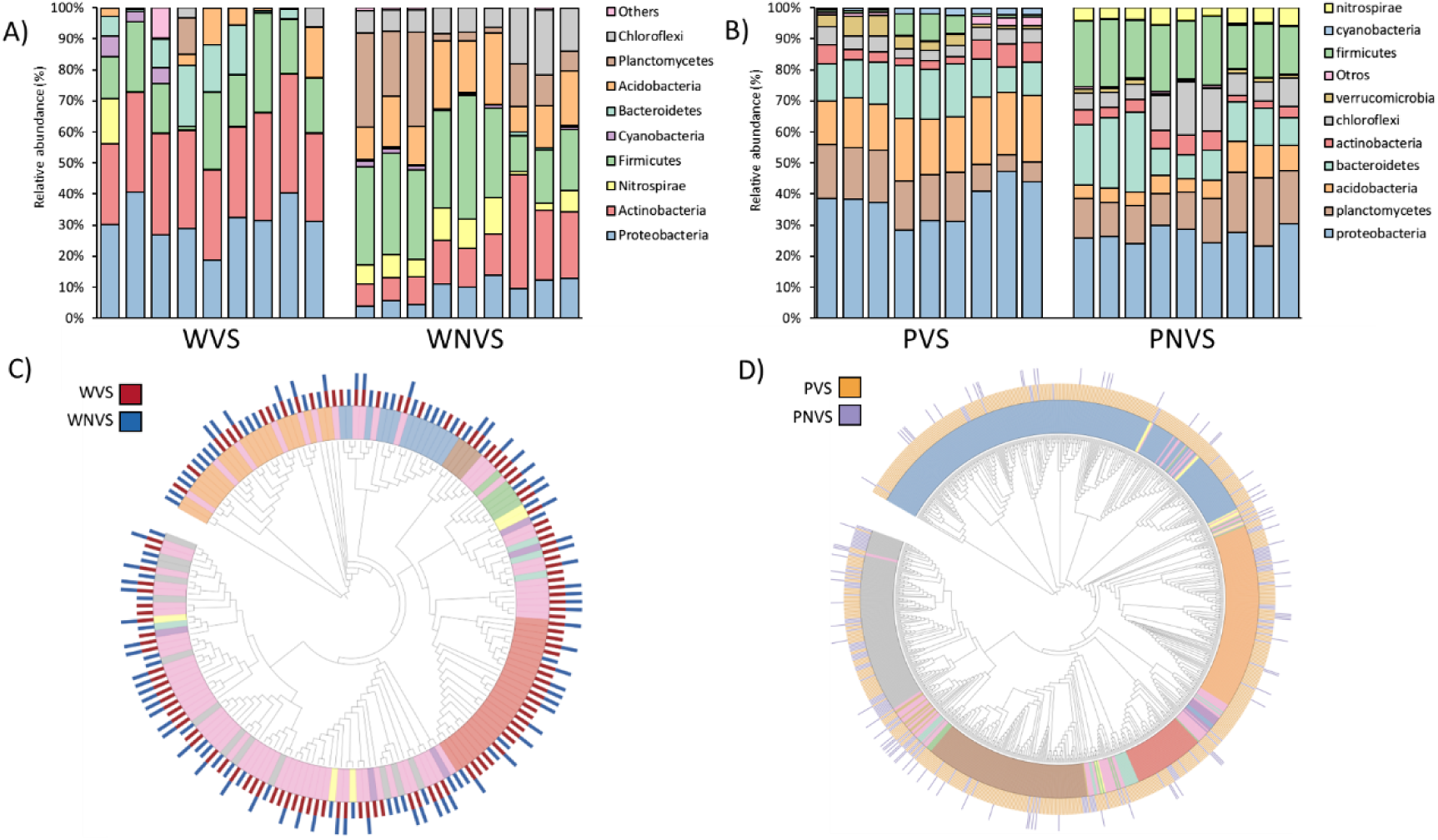
Taxonomic analysis of the bacterial community. A & B) Relative abundance of members of the microbial community from soils surrounding Willow (A) and Pine (B). C & D) Phylogenetic Tree of members of the microbial community from soils surrounding Willow (C) and Pine (D). Tree colors illustrate the phyla of each ASV and external column colors indicate the sampling stie of each ASV

Similar differences were observed between the pine sectors. PVS was predominantly represented by *Proteobacteria* and *Acidobacteria*, with 37.4% and 21.5% relative abundance, respectively. In comparison, PNVS had *Proteobacteria* (26.9%) and *Firmicutes* (19%) as the most abundant phyla. Although both sectors shared the most abundant phylum, PVS had a higher representation than PNVS (37.4% vs. 26.9%). Similarly, PVS had 3% abundance of *Firmicutes*, while PNVS had 6.1% *Acidobacteria*, highlighting the influence of pine on the abundance of different ASVs in the microbial community. To complement these observations, dendrograms were constructed to relate ASVs from WVS and WNVS (Figure 2C) and PVS and PNVS (Figure 2D). The results show that WVS and WNVS share a higher percentage of ASVs compared to PVS and PNVS, indicating that different sectors of the Cauquenes tailing have distinct soil microbiota compositions and that the impact of trees on these communities may depend on the tree species. Differences were also observed in the most abundant ASVs between WVS and PVS, particularly with *Actinobacteria* and *Planctomycetes*, being more abundant in WVS and PVS, respectively. To address this hypothesis, the effect of both plants on the soil’s environmental variables was analyzed. A canonical correspondence analysis (CCA) (Figure 3) showed significant differences in soil composition in terms of microbiota, with WNVS and PNVS clustering together, correlating directly with Cu, Mn, Fe, and Na. In contrast, the microbiota in WVS correlated with K, Ca, and P, a phenotype not observed in PVS, whose microbiota did not correlate with any studied variables, supporting the hypothesis that each tree species has a distinct effect on its environment.

**Figure 3.**
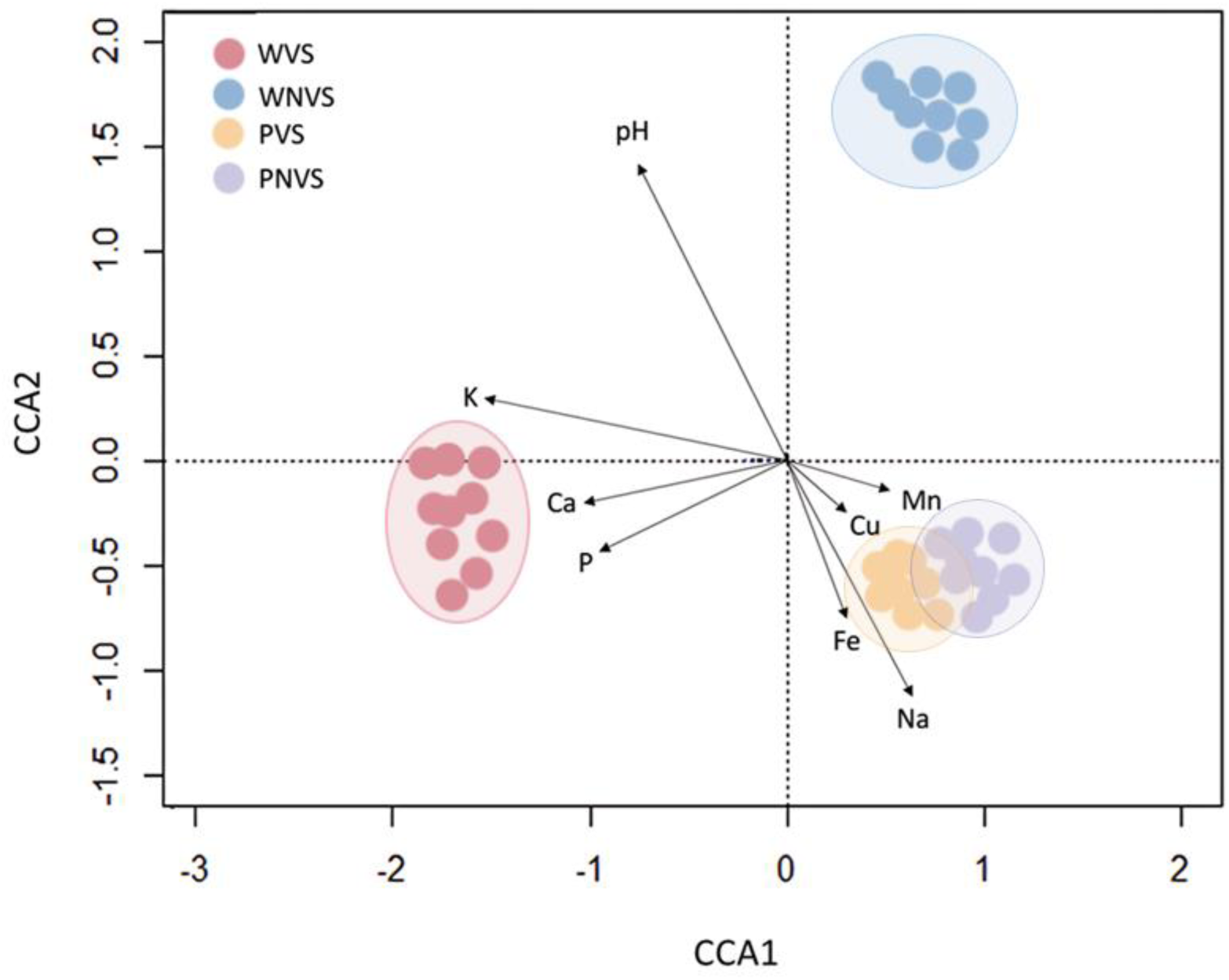
CCA diagram representing the relationship between tailings bacterial communities and their abiotic variables. The black vectors represent the direction of the variables that could explain the distribution between the points.

### 3.3 Co-occurrence Networks

To assess the interactions within the microbiota of each site, co-occurrence network models were constructed and analyzed for the four tailing sites (Figure 4A-B). First, the resulting networks exhibited distinct topological properties across the sites (Table 3). The WVS network consisted of 175 nodes and 1170 edges, while WNVS had 266 nodes and 2608 edges. The PVS network, being the most complex, contained 1360 nodes and 55593 edges. In comparison, the PNVS network had 371 nodes and 7440 edges. The WVS network demonstrated a high clustering coefficient and a high percentage of positive edges and high modularity, implying a stable community with a strong potential for cooperation through the formation of distinct functional niches (Dai et al., 2024). In contrast, the WNVS network had lower values for these metrics, suggesting that the presence of willow enhances the formation of a cohesive and cooperative community among the ASVs in the soil.

**Table 3.** Topological parameters of co-occurrence networks of each site. WVS: Willow-Vegetated Soil; WNVS: Willow Non-Vegetated Soil; PVS: Pine-Vegetated Soil; PNVS: Pine Non-Vegetated Soil.

| **Metric** | **WVS** | **WNVS** | **PVS** | **PNVS** |
| --- | --- | --- | --- | --- |
| Nodes | 175 | 266 | 1360 | 371 |
| Edges | 1170 | 2608 | 55593 | 7440 |
| Clustering coefficient | 0.892 | 0.573 | 0.532 | 0.621 |
| Modularity | 0.848 | 0.573 | 0.213 | 0.085 |
| Positive edge percentage | 97.7 | 58.2 | 56.1 | 51.9 |
| Edge density | 0.029 | 0.04 | 0.06 | 0.064 |

**Figure 4.**
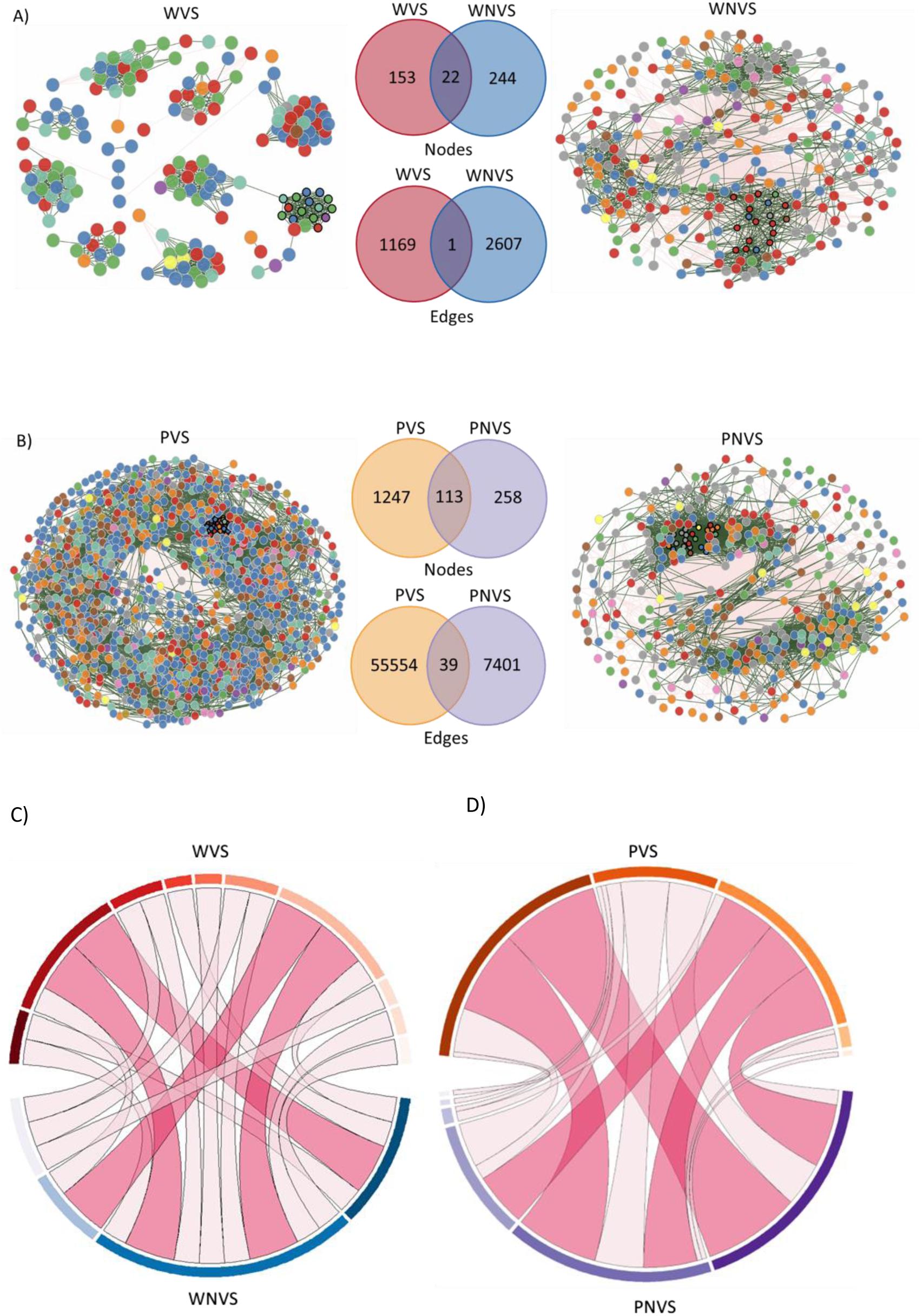
Microbial co-occurrence networks. A) Willow vegetated and non-vegetated soil networks, including common nodes and edges. B) Pine vegetated and non-vegetated soil networks, including common nodes and edges. The four networks were constructed based on the abundance data from each site. Bold border circles represent the hub nodes of each network. In the networks: green lines indicate positive interactions, while red lines indicate negative interactions. For common nodes and edges: warm colors (red and orange) represent those from vegetated soils, and cold colors (blue and purple) represent those from non-vegetated soils. C & D) Circos plots illustrating the modular structure and connectivity of bacterial communities in vegetated and non-vegetated soils. Warm colors (red and orange) represent modules from vegetated soils (WVS and PVS), while cool colors (blue and purple) represent modules from non-vegetated soils (WNVS and PNVS). The curved links between sections of the plot indicate shared nodes across modules and between soils

For the other sites, PVS and PNVS exhibited similar clustering coefficients and positive edge percentages, although PVS displayed higher modularity. This suggests that while the overall cooperation and interactions were similar, pine trees may influence the formation of functional groups within its network. Interestingly, WVS and WNVS shared 22 common nodes (Figure 4A), indicating a small number of shared nodes across these networks. In contrast, PVS and PNVS shared 113 common nodes (Figure 4B), representing 50% of the nodes in PNVS but only 9% of the total nodes in PVS. All networks shared a low number of edges (Figures 4F and 4H). These results indicate that the interactions between nodes differ across these networks, suggesting particular bacterial relationships between members of each specific site inside the Cauquenes tailings, relationships highly affected by the presence of pine or willow plants.

To determine the relationships between shared ASVs in each network, two Circos plots were constructed, illustrating the WVS (red) and WNVS (blue) networks (Figure 4C) and the PVS (orange) and PNVS (purple) networks (Figure 4D). Both Circos plots demonstrate that the modularity among shared ASVs is influenced by the presence of vegetation, with different modules forming between non-vegetated and vegetated soils. Modules are groups of bacterial taxa (ASVs) that are more densely interconnected within themselves than with other groups, reflecting functional or ecological interactions (Li et al., 2021). These findings corroborate previous results (Mandakovic et al., 2023; X. Sun et al., 2018), highlighting not only the impact of plants on the tailing environment but also the plant-specific nature of these effects.

### 3.4 Functional Metabolic Prediction of the Tailing Microbiota

To robustly describe a microbial community, it is essential to study interactions, functionality, and assembly. First, the PICRUSt 2 tool was used to predict the functions of the microbiota in the four sectors studied. In general terms, basal metabolism was similar across all sites, with only minor differences in aromatic compound degradation, carbohydrate degradation, fatty acid and lipid degradation, and secondary metabolite synthesis. However, these minor differences allowed for the classification of two groups (Figure 5A): sites without tree growth (WNVS and PNVS) and sites with tree growth (WVS and PVS), suggesting that tree growth also influences bacterial community functions, independent of species. To complement the findings from PICRUSt 2, which focuses on global metabolic processes, the consumption of 31 different carbon sources was determined using Biolog EcoPlates. Clustering based on consumed metabolites once again separates the communities into two major groups (Figure 5B). As expected, the communities at WNVS and PNVS showed low utilization and metabolism of the substrates tested, with near-detection limit intensity. In contrast, the WVS community utilized a greater number of substrates and showed higher metabolism than sites without tree growth. Finally, the PVS community exhibited the highest number of metabolized substrates, and the greatest metabolism compared to other sites. These differences indicate that tree presence makes the bacterial community more active in carbon source utilization, with species-specific effects.

**Figure 5.**
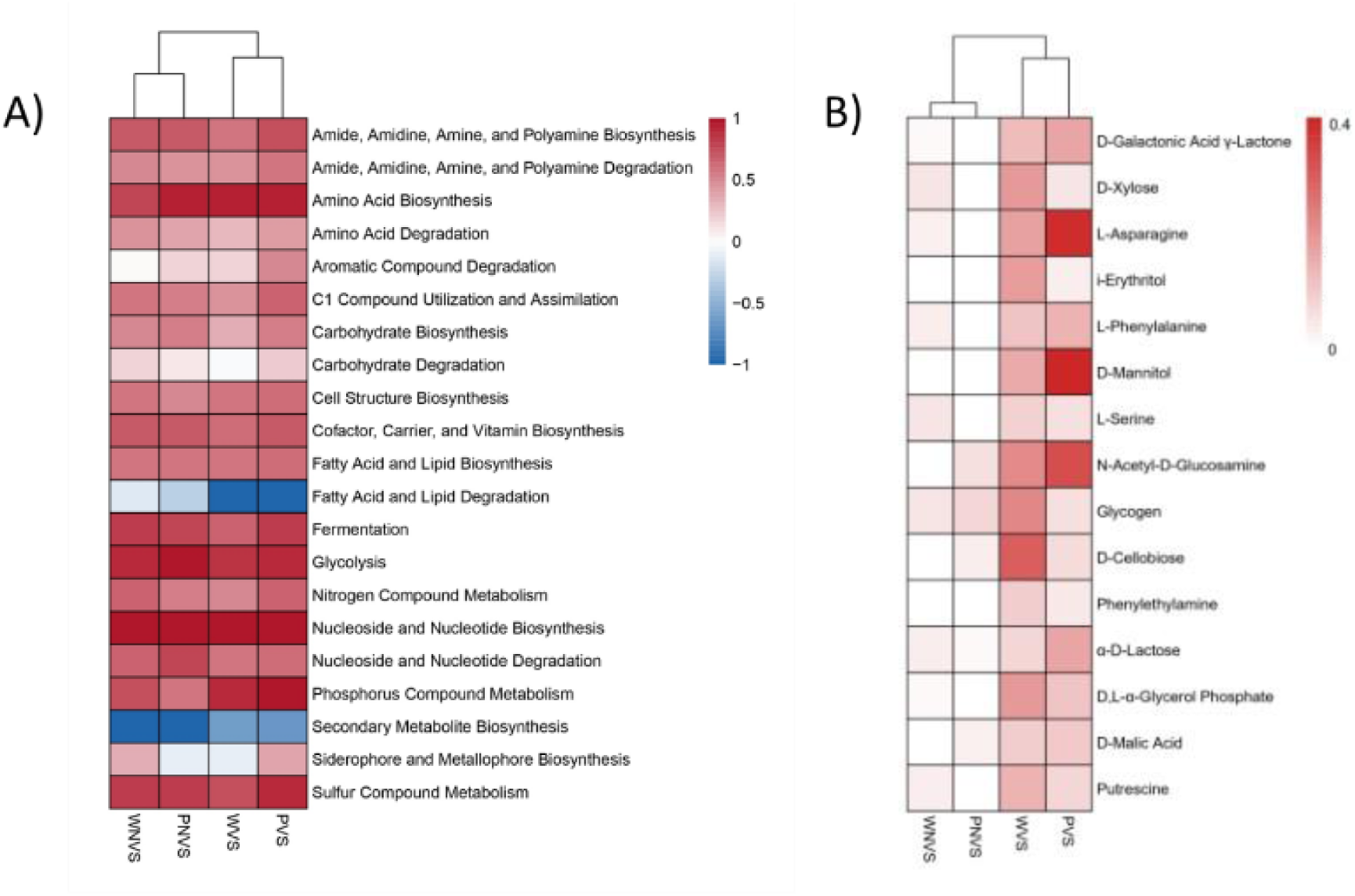
Metabolic prediction and carbon source consumption of bacterial communities from each site. A) Heatmap illustrating the predicted metabolic functions of bacterial communities at each site, based on PICRUSt2 analysis. The color gradient ranges from red (indicating higher predicted metabolic activity) to blue (indicating lower predicted metabolic activity). B) Carbon source utilization profiles of bacterial communities at each site, visualized as a heatmap based on Average Well Color Development (AWCD) from Biolog EcoPlate analysis. The intensity of red indicates the level of carbon source consumption, with darker red denoting higher utilization rates.

ΒNTI was calculated to assess whether the assembly of the microbial community in each sampling site is governed by deterministic or stochastic processes (Figure 6). |ΒNTI| values for sites with tree growth (WVS and PVS) were > 2, indicating that their bacterial communities are assembled by deterministic processes and have a homogenous selection (ΒNTI < -2). On the other hand, sites without tree growth (WNVS and PNVS), showed |ΒNTI| values < 2, indicating that stochastic processes, such as ecological drift or dispersal, are playing a significant role in the community assembly. These results indicate that tree growth played a key role in shaping the microbial community.

**Figure 6.**
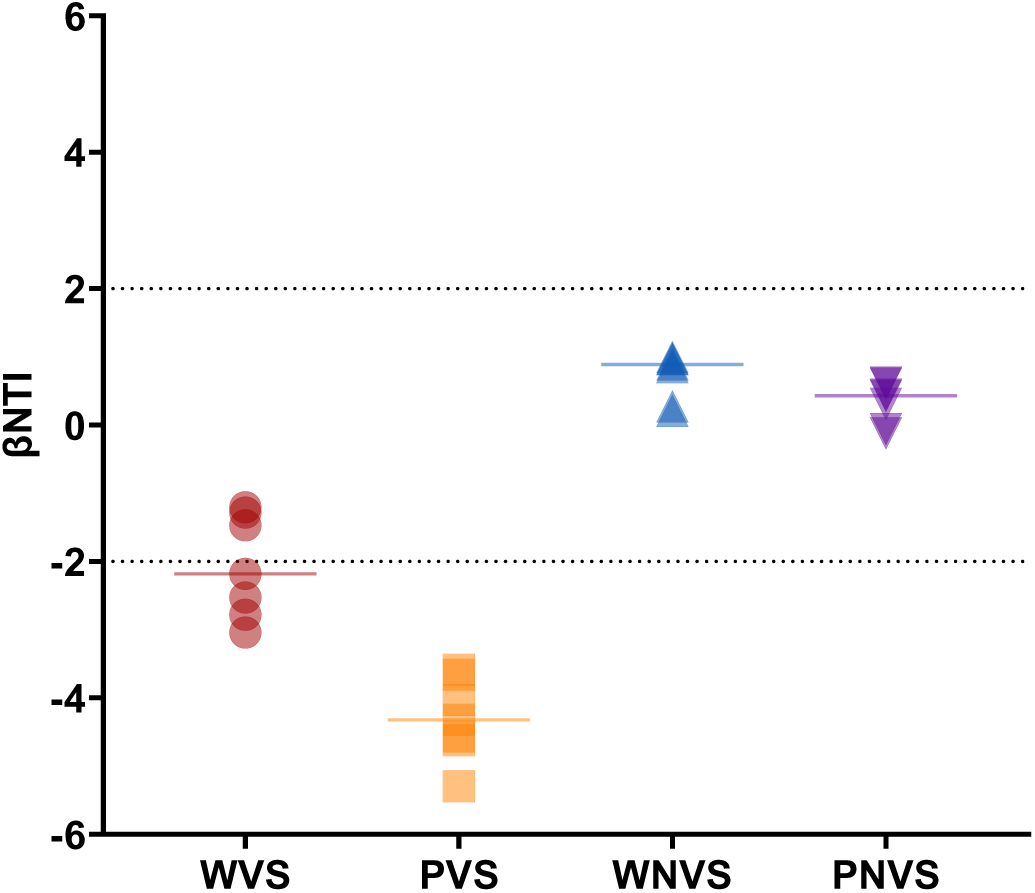
βNTI distribution highlighting the assembly mechanisms of the bacterial community from vegetated and non-vegetated soils. Dashed black lines (ΒNTI > 2 and ΒNTI < 2) represent the 95% confidence interval of the standard normal distribution.

## 4. Discussion

In this study, we investigated the effects of willow and pine tree growth in Cauquenes tailings. We focused on analyzing the concentration of elements, microbial communities and interactions, and metabolic capabilities. Our results highlight different patterns of metal accumulation, microbial diversity and metabolic capabilities, depending on the proximity and type of tree.

### 4.1 Phytoremediation Potential of Willow and Pine in a Copper Mine Tailing

(Mendez & Maier, 2008; Salt et al., 1995) Tailings, in general, have low content of nutrients and high concentration of potential toxic metals, which results in an unfavorable environment for most organisms (Pérez et al., 2021). Considering this, there’s numerous evidence of plant growth in mining tailings (Becerra-Castro et al., 2012; Gagnon et al., 2020; Gazitúa et al., 2021), including in Cauquenes (Aponte et al., 2023). In a previous work, we identified different zones within Cauquenes tailing, varying in metal concentrations and pH values (Galvez et al., 2022) In relation to this, Cu, Zn and Na in non-vegetated soils were higher than New tailings soil but lower than Old tailings soil. On the other hand, they were similar to soils studied in (Aponte et al., 2023), highlighting that Cauquenes tailing has different soil conditions in its expanse. Plant growth usually results in a modification of the concentration of chemical elements in the soil, thanks to absorption and accumulation in different tissues of the plant (Yan et al., 2020). Regarding this, in vegetated soils we found lower concentrations of potential toxic elements for plant development, such as Cu, Zn, and Na. These lower concentrations may be attributed to adsorption by plant litter (Wu et al., 2018), humic substances (Kulikowska et al., 2015), complex formation of metals with root exudates (Kim et al., 2010), or uptake by the roots of the plants (Y. Liu et al., 2023). There’s evidence of absorption of these elements by willows: A 2009 study by Kacálková *et al*. found that willows in contaminated soils accumulated high levels of Zn and some Cu in their leaves (Kacálková et al., 2009). Similarly, a 2011 study showed willows at a Japanese mining site absorbed and accumulated Zn in their leaves (Harada et al., 2011). In a similar vein, pines had also been shown to absorb these metals (Asensio et al., 2013; Martinez-Oró et al., 2017; Parraga-Aguado et al., 2014). To test this hypothesis, we took samples from roots and leaves from willow and pine (WR, WL, PR, PL, respectively). Broadly speaking, all elements were higher in plant tissues than in the soils. Specifically, Cu was higher in WR than in WL, indicating that this element may not be distributed to aerial plant tissues. This could indicate a phytostabilization potential for Cu (Mendez & Maier, 2008; Salt et al., 1995). On the other hand, Zn and Na were higher in WL than WR, indicating that Willow has a potential for phytoextraction of these elements (Suman et al., 2018). In Pine tissues, we found that Cu and Zn accumulate on leaves rather than roots, indicating a phytoextraction of these elements. Overall, our results suggest that both plant species could be useful for future phytoremediation of this Tailing.

### 4.2 Tree-Dependent Reshaping of Soil Bacterial Communities in Copper Mine Tailings

The conditions of the tailings not only affect plant development, but also microbial growth. Since plant growth reduced the concentration of toxic elements in the soil, we set out to analyze its effects on the bacterial composition within vegetated and non-vegetated soils. Of all the soils, PVS was the one that had the most taxonomically rich site, showing the presence of 1199 ASVs. In comparison, PNVS had 435 ASVs, almost 3 times less than its vegetated soil. This could be attributed to the ability of plants to modulate its microbiome via the recruitment of beneficial bacteria (Kwak et al., 2018). Although this recruitment of beneficial bacteria could explain the differences in ASV composition, it alone does not fully account for why PVS had significantly more ASVs than PNVS. This phenomenon could be attributed to the way that pine roots behave in this environment, being long and superficial, they could be recruiting bacteria across long patches of land. This could also be explained by airborne bacterial recruitment via the phyllosphere (Santoyo, 2022), and thanks to deposition of pine litter, these bacteria could enter the soil thanks to interactions with pine roots. On the other hand, WVS had less ASVs than WNVS, an expected result, since it’s known that plants can recruit their own microbiome and adapt it to better survive in harsh environmental conditions (Monohon et al., 2021; Rolfe et al., 2019).

To further explore the effects of willow and pine vegetation, we analyzed the relative abundance of bacterial phyla in each sampled site. In WVS, the most abundant ASVs were Actinobacteria (31.3%) and Proteobacteria (31.3%), while in WNVS the most abundant ASVs belonged to Firmicutes (26.8%) phylum. This could be explained by the neutral pH that WVS had, since it has been reported that neutral pH favors the growth of *Actinobacteria* and *Proteobacteria*, while acidic pH favors *Firmicutes* growth (D. Qi et al., 2018; C. Wang et al., 2019). Another explanation could be that Willow is secreting exudates that recruit bacteria belonging to those phyla. While there’s numerous reports that Willow growth is correlated with higher abundance of *Proteobacteria*, with *Actinobacteria* is not the case (Koczorski et al., 2022; Tardif et al., 2016; G. Wang et al., 2021). The higher abundance of *Actinobacteria* in our case may be due to the microbial recruitment of this phyla from WNVS, in which it had approximately 10% of the total abundance. There are reports that mention that bacteria belonging to Actinobacteria have resistance to heavy metals (Cu included) (Presentato et al., 2020; Schmidt et al., 2005), potential for remediation of heavy metal polluted soils (Benimeli et al., 2011; Kannabiran, 2017), and they have members with potential PGP traits (Boukhatem et al., 2022; Hamedi & Mohammadipanah, 2015). All these reasons combined make *Actinobacteria* a suitable candidate for recruitment from Willow trees. More importantly, the study published by (Touceda-González et al., 2017) found that the presence of *Salix viminalis*, after compost application in a Cu-mine tailing in Spain provoked an increased abundance of *Actinobacteria*, along α-*Proteobacteria* and β-*Proteobacteria*, supporting the findings in our study.

In the case of pine sites, PVS ASVs were predominantly Proteobacteria (37.4%) and Acidobacteria (21.5%). PNVS bacterial abundance was mostly represented by Proteobacteria (26.9%) and Firmicutes (19%). As with WVS, this higher abundance of Proteobacteria could be explained by the pH in this site. Similarly, various studies have shown that Pine growth could be correlated with more abundance of Proteobacteria in soil (Chow et al., 2002; Padda et al., 2022; Proença et al., 2017), and of Acidobacteria (Y. Ma et al., 2020; Zheng et al., 2020). While we didn’t found reports of PGPB from Acidobacteria, its known that members of that phylum can help restore soil functionality by participating in carbon and nitrogen cycles, organic matter decomposition, and soil structure enhancement by secreting exopolysaccharides (Garcia-Villaraco et al., 2024; Kalam et al., 2020), so while these bacteria may not be interacting directly with pine trees, they can promote its growth by restoring soil health.

### 4.3 Willow and Pine Modulate Soil Bacterial Interactions in a Copper Mine Tailing

To further explore the effects of tree growth in the bacterial community structure and arrangements, we studied ASVs interactions by constructing and analyzing co-occurrence interaction networks in each study site (Mandakovic et al., 2018). Regarding networks of WVS and WNVS sites, we found that the vegetated soil had less nodes and edges, indicating that its community has less members and interactions. This could mean that Willows could be specifically selecting its microbiome, leaving beneficial bacteria and eliminating pathogenic ones, shaping their interactions, making them specialized in supporting the growth of the tree. These findings align with previously published studies (L. W. Mendes et al., 2014; R. Mendes et al., 2013), including those done in mine tailings (Romero et al., 2021) and they’re also supported by the other topological parameters in the network, such as its higher modularity, clustering coefficient and positive edge percentage (Fabbrini et al., 2023; C. Liu et al., 2023). In contrast, PVS network had significantly more nodes and edges than its non-vegetated counterpart. While we expected recruitment and selection of beneficial ASVs, reflected in a shift in nodes such as the one seen in WVS network, we didn’t expect a recruitment of almost ten times more ASVs. Previously, it has been reported that revegetation efforts in an acid mine tailings pond provoked the recruitment of more ASVs in the network of soils surrounding plant growth, and that this recruitment is dependent on the type of revegetation and plant species. One type of revegetation, called Capped Revegetation, is carried out by applying a new soil cover from other ecosystems and then planting the trees (Zhou et al., 2020). Also, the roots of these trees are superficial and cover extended distances. These roots could be recruiting airborne microorganisms and integrating them into the network. In a similar vein, this network showed higher modularity than its non-vegetated counterpart, indicating formation of microbial modules which could specialize in plant-growth promoting parameters, supporting pine growth. This shift in modularity is supported by our Circos plots, which show how the common nodes in WVS / WNVS, and PVS / PNVS have different arrangements in the presence of the trees. Overall, our results show that the microbial communities are affected by tree growth in Cauquenes tailings, not only in its relative abundance but also in their interactions between each member, and these differences are dependent on the type of tree.

### 4.4 Shift in Metabolic Profiles of Soil Bacteria depends on the Presence of Willow and Pine

Despite the differences between community structure between each study site, it’s possible that the function that they play at each study site might be similar, we set out to study the metabolic profile of each bacterial community.

PICRUSt 2 was used to predict the metabolic functions of the community at each study site (Zhao et al., 2021). Overall, the most abundant metabolic groups were related to amino acid synthesis, fermentation, glycolysis, nucleoside and nucleotide synthesis, and sulfur compound metabolism. These functions have been previously reported to be enriched in tungsten, Cu, and cadmium mine tailings (Chung et al., 2019; Zhao et al., 2021; Zhu et al., 2022). Glycolysis could produce pyruvate and subsequently, fermented to produce organic acids (Newsome & Falagán, 2021). These organic acids can enhance the mobility of Cu ions by lowering the pH of the environment, increasing its bioavailiability to plants and assisting in Cu remediation (Pérez-Esteban et al., 2013). Nucleoside and nucleotide synthesis could be related to increased production of ATP, which is a core molecule in the function of P-Type ATPases, related to numerous heavy-metal resistance proteins, such as CopA or ZntA (Palmgren, 2023). Lastly, sulfur compound metabolism could be related to the presence of important bacterial species in the sulfur cycle. (Zhao et al., 2021). A clustering of these results was performed to further explore the effects of vegetation in the metabolism of the community. Our results grouped together the vegetated sites and the non-vegetated sites, indicating that, at this level, the difference between the metabolic profiles depends on the presence or absence of vegetation, and not the type of tree. The main differences were seen at Fatty Acid and Lipid Degradation (higher in WNVS and PNVS), and at Secondary Metabolite Biosynthesis (higher in WVS and PVS). Microbial communities in WNVS and PNVS, due to the lower organic matter content, could shift their metabolism to use fatty acids and lipids as a carbon source (Larkin, 2003), in comparison with WVS and PVS which could be utilizing plant litter or root exudates as carbon sources (Kramer & Gleixner, 2008). The presence of these exudates could also be related to the Secondary Metabolite Biosynthesis in WVS and PVS (Benizri et al., 1998).

The Biolog EcoPlate system can be used to further explore the metabolism of the microbial communities, focusing on the metabolism of 31 different carbon sources (Feigl et al., 2017). Overall, non-vegetated sites showed a poor utilization of these carbon sources, which correlates to previous studies (Lebrun et al., 2021; Ultra et al., 2013). These sites primarily metabolized Tween 40 and Tween 80, which are known tensoactives with a lipid-like structure, which could be related to the higher lipid degradation shown in PICRUSt 2 analysis. On the other hand, WVS and PVS showed a higher quantity of carbon sources metabolism, which could be related to the adaptations that the trees produced in the microbial community. Of note, the type and quantity of carbon sources utilization differed between WVS and PVS. Specifically, PVS showed a wider variety of carbon sources metabolism than WVS, which could be related to the more taxonomically rich community in these sites.

βNTI is commonly used to predict the effects of deterministic and stochastic processes on microbial communities (Su et al., 2022). In this work, bacterial communities from vegetated sites (WVS and PVS) had ΒNTI values associated with a deterministic assembly. This was an expected result, since it has been previously determined that bacterial communities’ assemblage from restorated soils are governed by deterministic processes (|βNTI| > 2) (W. Gao et al., 2020; Long et al., 2024; K. Wang et al., 2022). This is also true in revegetated soils from a mining environment (Yang et al., 2017). PiCRUST 2 and Biolog EcoPlate results also correlate with the βNTI value, since communities from vegetated soils group together in the dendrograms. On the other hand, non-vegetated soils (WNVS and PNVS) had βNTI values correlated with stochastic processes (|βNTI| < 2), which include dispersal and ecological drift (Xia et al., 2023). We also expected this result, since reports from Cu mining soils also showed that the assembly of its bacterial community is governed by stochastic processes (B. Liu et al., 2022; X. Ma et al., 2023). These findings collectively highlight that vegetation could drive a shift in bacterial community assembly processes from stochastic to deterministic in mining-impacted soils.

## 5. Conclusions

Mine tailings are an extreme environment that is generally unfavorable for plant growth and establishment. However, in the Cauquenes tailings, there is growth of trees. Given the importance of bacterial and plant interactions, this study focused on analyzing the effects of the presence of willow and pine trees on heavy metal concentrations and bacterial abundance and interactions. The presence of both trees resulted in lower heavy metal concentrations in the soil, while accumulating them in their roots and leaves, indicating that these trees have potential to be used as bioremediators for Cu in these terrains. Both trees also caused a shift in bacterial abundance compared to the non-vegetated sites, increasing the abundance of Proteobacteria, Actinobacteria, and Acidobacteria. Bacterial networks in the vegetated sites also exhibited higher modularity, suggesting the formation of functional groups. The bacterial communities in the vegetated sites showed higher metabolic activity, carbon source consumption, and were assembled by deterministic processes. All these characteristics could be associated with an improved environment for tree growth. Our results illustrate the importance of plants in mitigating contaminated mining environments, not only by reducing the concentration of heavy metals but also by promoting a more active bacterial community.

## CRediT authorship contribution statement

**Jaime Ortega**: Methodology, Formal analysis, Investigation, Writing – original draft, Writing – review & editing, Visualization. **Gabriel Gálvez**: Methodology, Formal analysis, Validation, Investigation, Writing – review & editing. **Gladis Serrano**: Methodology, Investigation. **Jorge Torres**: Visualization, Validation. **Victor Aliaga-Tobar**: Formal analysis, Validation. **Emilio Vilches:** Visualization, Validation. **Angélica Reyes:** Visualization, Validation. **Vinicius Maracaja-Coutinho:** Formal analysis, Validation. **Valentina Parra**: Formal analysis, Visualization, Validation. **Alex Di Genova:** Formal analysis, Validation. **Lorena Pizarro:** Conceptualization, Visualization, Validation. **Mauricio Latorre**: Conceptualization, Writing – review & editing, Visualization, Supervision, Resources, Project administration, Funding acquisition.

## Declaration of competing interest

The authors declare that they do not have any competing financial interests of personal relationships that could influence the results of this research

## Acknowledgments

This work was supported by Proyecto ANILLO regular ANID ACT210004, Center for Mathematical Modeling, Apoyo a Centros de Excelencia ACE210010; ANID Millennium Science Initiative Program ICN2021_044; Fondo Basal FB210005; ANID FONDECYT 1190742 and 1230194, Fondo Interdisciplinario y Proyecto Núcleo UOH, Proyecto Postdoctoral ANID 3220080, Becas ANID 21211367 and 21220593. We appreciate the contribution of Minera Valle Central (Chile), SEREMI de Minería Región de O’Higgins and CODELCO.

## Notes

### Competing Interest Statement

The authors have declared no competing interest.

